# A global perspective on bioinformatics training needs

**DOI:** 10.1101/098996

**Authors:** Michelle D. Brazas, Cath Brooksbank, Rafael C. Jimenez, Sarah Blackford, Patricia M Palagi, Javier De Las Rivas, B.F. Francis Ouellette, Judit Kumuthini, Eija Korpelainen, Fran Lewitter, Celia W.G. van Gelder, Nicola Mulder, Manuel Corpas, Maria Victoria Schneider, Tin Wee Tan, Dave Clements, Angela Davies, Teresa K. Attwood

## Abstract

In the last decade, life-science research has become increasingly data-intensive and computational. Nevertheless, basic bioinformatics and data stewardship are still only rarely taught in life-science degree programmes, creating a widening skills gap that spans educational levels and career roles. To better understand this situation, we ran surveys to determine how the skills dearth is affecting the need for bioinformatics training worldwide. Perhaps unsurprisingly, we found that respondents wanted more short courses to help boost their expertise and confidence in data analysis and interpretation. However, it was evident that most respondents appreciated their need for training only after designing their experiments and collecting their data. This is clearly rather late in the research workflow, and suboptimal from a training perspective, as skills acquired to address a specific need at a particular time are seldom retained, engendering a cycle of low confidence in trainees. To ensure that such skill gaps do not continue to create barriers to the progress of research, we argue that universities should strive to bring their life-science curricula into the digital-data era. Meanwhile, the demand for point-of-need training in bioinformatics and data stewardship will grow. While this situation persists, international groups like GOBLET are increasing their efforts to enlarge the community of trainers and quench the global thirst for bioinformatics training.

## Introduction

Over the past decade, anecdotal evidence of a bioinformatics skills gap amongst life scientists has been growing (Abeln *et al.*, 2013;Brass, 2000; MacLean & Miles, 1999; Schneider & Jungck, 2013). In 2013, this prompted the Society for Experimental Biology (SEB) to work with the Global Organisation for Bioinformatics Learning, Education and Training (GOBLET;Attwood *et al.*, 2015) to run a brief survey to capture a snapshot of training needs within the SEB community: the questionnaire focused on the ways in which bioinformatics training and education had been received; the level of confidence in using bioinformatics databases, software and command-line tools; how respondents wanted bioinformatics training to be delivered; and, in particular, what skills were needed or most demanded. The survey garnered ~200 responses.

Trends in the results were sufficiently interesting to prompt GOBLET to push the survey out more widely through its partner organisations (throughout Europe, North America, Africa, Asia and Australia), to try to establish whether the initial findings were general across global communities of life- and computational scientists. Outcomes of this second survey, which attracted ~500 responses, were discussed at a bioinformatics workshop held alongside the SEB annual meeting in July 2014 (SEB/GOBLET Bioinformatics Workshop, 2014). Workshop participants were invited to reflect on whether the survey trends resonated with them, whether there were gaps in the survey results, and what they felt providers of bioinformatics education and training could do to help address the identified gaps (*e.g.*, through the provision of short professional courses, summer schools, workshops, or online courses and training modules).

The results of these and other GOBLET member surveys, together with feedback from the workshop, were presented and discussed during the GOBLET annual meeting held in Toronto in November 2014. While the surveys do not reflect a rigorous and complete analysis of global bioinformatics training needs, they nevertheless provide some initial perspectives on the current status of training gaps and needs across GOBLET’s national and regional member organisations. From this, it is clear that the need for bioinformatics training is both real and urgent, and requires worldwide solutions.

The driver behind the need for bioinformatics training is the landscape of life-science research today - it is voluminous, complex and integrative, and is increasingly computational. Bioinformatics skills have become intrinsic to life-science research, particularly to ‘omic’-based research (proteomics, genomics, metagenomics, *etc.*). While rudimentary programming, use of bioinformatics tools and databases, and statistical principles are beginning to appear in some undergraduate (UG) life-science degree curricula (Goodman & Dekhtyar, 2014; Libeskind-Hadas & Bush, 2013; Rubinstein & Chor, 2014), basic skills in data analysis and interpretation, and especially in data management, are still taught relatively rarely within traditional UG programmes, even though such skills are essential to most research projects today. Many students thus progress to postgraduate (PG) life-science degrees without adequate foundations in computational science. Hence, there is still a profound gap between the theory learned at degree level and the real-life practice of research data analysis.

This observation resonates with the outcomes of a 2014 survey run by the Association of the British Pharmaceutical Industry (ABPI): respondents from a range of pharmaceutical companies, contract research organisations, and small and medium enterprises identified major skills gaps in mathematical and computational areas, leading to significant problems with both the number and quality of candidates available for recruitment (ABPI, 2015). The dearth of computational and statistical skills is not just an issue for life scientists, but has been recognised in graduates in many scientific, medical and business-related disciplines, providing the impetus for the recent emergence of new university programmes and training courses in ‘generic’ data science.

Of course, not all scientists involved in research need to become bioinformaticians, but acquiring at least a minimum level of bioinformatics understanding (Tan *et al.*, 2009) can help life- and computational scientists to communicate and interact with one another more effectively (whether in discussions about experimental design or particular analyses, or about specific technical requirements), as well as improve critical thinking about their research findings. Even experienced bioinformaticians - many of whom are self-taught - need to garner new skills to keep up with evolving, leading-edge technologies, or to reinforce knowledge they gained at some point in the past. Although individual factors may differ (career development, change in job responsibilities, change in research project, new technology, *etc.*), bioinformatics skill gaps clearly impede advancement, and are ultimately fuelling a global thirst for bioinformatics education and training.

## Audiences seeking bioinformatics training are broad

The survey was kept brief, as we aimed to gain a high-level picture and wanted to encourage participation (see Table 1, Supplementary Data for the full set of questions). Examining who required bioinformatics training revealed a broad and expansive audience, not confined to the wet-lab researcher. From the results, those seeking bioinformatics training could be seen to span the full science career trajectory, and a range of job titles, an observation not limited to any particular geographical location.

**Table 1:** Questions used to survey the bioinformatics training needs of the SEB and GOBLET communities.

|  | Survey questions |
| --- | --- |
| 1 | What is your position/job title? |
| 2 | What is your main research discipline? |
| 3 | In which research institute/university/organisation and in which country? |
| 4 | Which bioinformatics databases, software tools, analysis packages and/or interpretation techniques do you currently use in your research and for what purpose? |
| 5 | How confident are you using bioinformatics databases, tools, techniques, *etc.*? |
| 6 | How have you acquired bioinformatics knowledge/skills (past and present)? |
| 7 | Which bioinformatics skills training would you most value (*e.g.*, database selection, software tools, analysing and interpreting data)? |
| 8 | How would you prefer training to be delivered to you? |
| 9 | Any other comments you would like to add? |

As Figure 1 illustrates, training needs were expressed by postgraduate students, through research fellows and technical staff, to senior professionals (including managers, group leaders and professors). The respondents originated from academia, healthcare and industry, cutting across a spectrum of scientific disciplines (including mathematics, computer science, biophysics, biochemistry, bioinformatics, biocuration, genomics, biomedical/plant/marine sciences, and so on) (data not shown).

**Figure 1:**
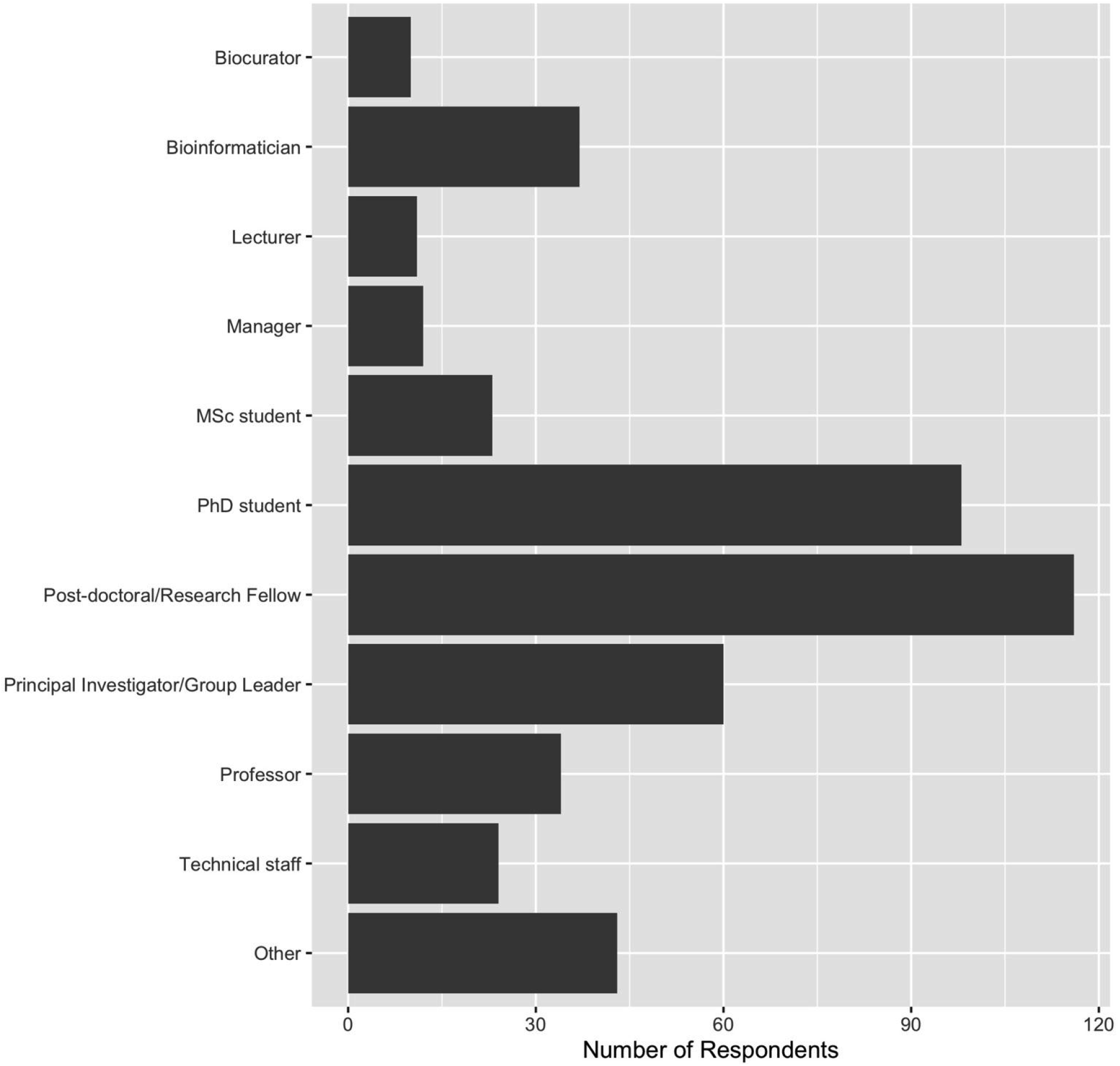
Chart showing respondents to the GOBLET survey in terms of their career stage/job title.

Given the new research avenues opened up by the advent of high-throughput platforms, and the impact these are having on the life sciences, these results are not surprising. The volumes of data being produced by such platforms necessitate the use of bioinformatics tools and resources, statistical analysis techniques and high-performance computing, bringing with them a growing demand for training.

While training needs differ according to individual academic backgrounds and job specifications (as discussed below), even computational biologists and bioinformaticians cannot be experts in all tools and techniques. For example, software and database versions, and their user interfaces, are in a continual flux as updates, upgrades and improvements are made. Moreover, analysis methods and approaches are also constantly evolving as particular bioinformatics fields (*e.g.*, Next-Generation Sequencing (NGS) data analysis) progress towards maturity and stability. These shifting sands require all users to have ongoing access to training in order to remain current in their particular fields.

Overall, the nature of the audiences requesting training showed similar trends regardless of the country or region surveyed. Indeed, similar surveys conducted by GOBLET member organisations and other anecdotal evidence concur that bioinformatics training is fundamental globally across all educational levels, career roles, stages and sectors in the life- and computational sciences.

## What are the bioinformatics training needs of these audiences?

Collectively, not only did the surveys show that the audiences requiring bioinformatics training are broad (Figure 1), they also highlighted that the training they require covers a wide variety of topics. Above all, however, the greatest need expressed by the majority of respondents was for skills in data/statistical analysis and interpretation. Interestingly, these needs mirror the requirements sought by bioinformatics core facility directors (Welch *et al.*, 2014). The ability to select and use databases, and to use and adapt software tools, was also considered important, as was some level of introductory programming, but these were given far less priority than data analysis.

## How have these audiences learned and how do they want to learn?

We looked both at how respondents had received bioinformatics training in the past, and at how they would prefer to acquire bioinformatics skills in the future. The feedback (summarised in Figure 2) showed that most respondents considered themselves to be self-taught, including consulting with, and taking advice from, colleagues and peers, and using online materials and resources; some had also enrolled in face-to-face professional training courses. A little more than 25% reported that they had acquired their skills via formal university education programmes at either UG or PG levels.

**Figure 2:**
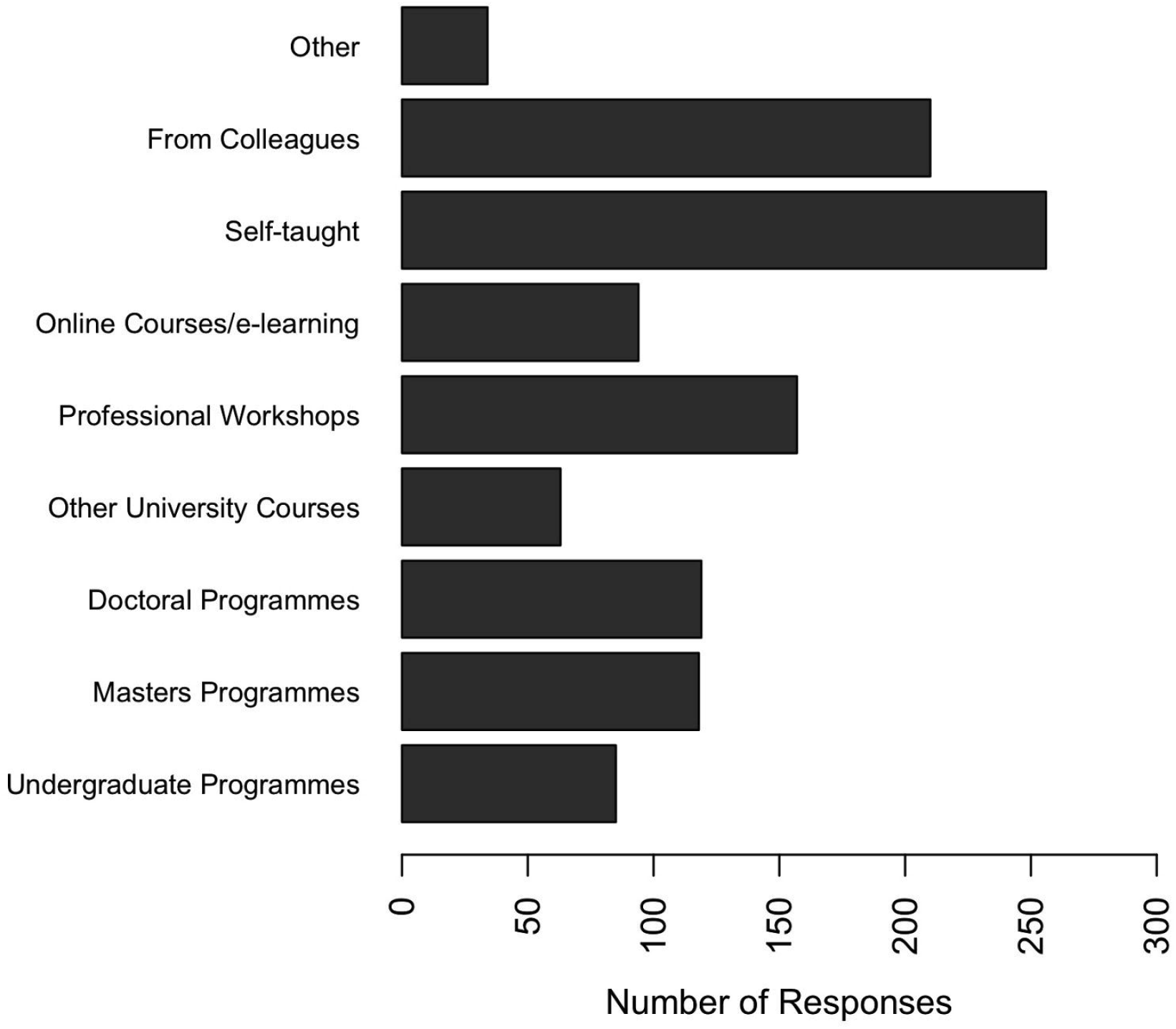
Chart showing how respondents to the GOBLET survey had acquired their bioinformatics skills.

In terms of how respondents would like to receive bioinformatics training in the future (Figure 3), most requested face-to-face workshops, whether delivered in the workplace or alongside conferences, or as stand-alone professional courses and summer schools. Many continued to favour access to online learning, several placing particular emphasis on the added value of tutor-supported options (e.g., free-text responses revealed, “*Online course ok if includes a good opportunity to discuss with teachers*,” and “*As an absolute beginner, it is very difficult to even begin to try to use command-line software and all previous courses have been too advanced for someone who has never used Linux OS. It would be helpful if there were opportunities to work through example data in small groups with a tutor*”). Similar findings have been reported previously (Lim *et al.*, 2003).

**Figure 3:**
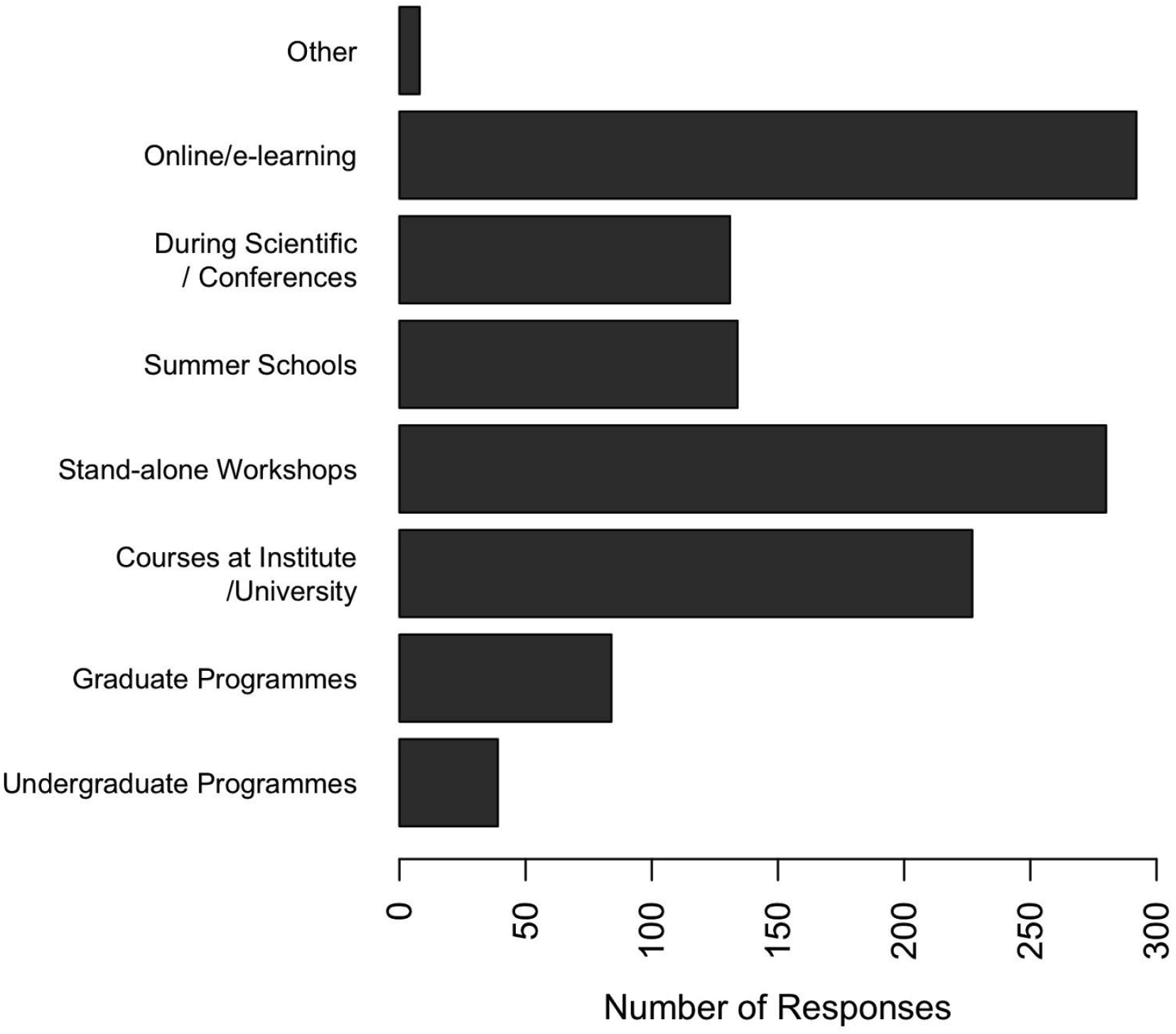
Chart showing how respondents to the GOBLET survey preferred training to be delivered.

UG and PG programmes (which are typically tutor-led) were noticeably less popular as future routes for bioinformatics training.

## How confident are these audiences with bioinformatics analyses, tools and resources?

Given how respondents reported having received bioinformatics training, and what training they considered they still needed, it was interesting to examine how confident they currently felt in using bioinformatics tools and resources, working at the command line and performing data analyses (see Supplementary Data, Figure 1).

Not surprisingly, bioinformaticians reported the greatest levels of confidence overall, particularly with regard to working at the command line. In general, however, the perceived confidence with data-analysis techniques was not conspicuously low. This is curious, given that the most sought-after skills expressed by respondents were in data analysis and interpretation, well beyond the needs they expressed for training with tools and resources.

## Understanding training needs and solutions

Identifying appropriate training for specific individuals typically depends on a number of factors, including 1) the reasons for seeking training; 2) the trainees’ background knowledge; 3) the format of training (including depth, level of difficulty and nature of training required); and 4) the time available for training. These factors (all of which will have influenced the survey responses) and their interdependencies must be considered in order to better understand the real training needs and solutions for specific audiences. We explore these factors briefly here.

### 1. Reasons for Seeking Training

The reasons for seeking bioinformatics training necessarily vary. Some individuals, for example, may need to improve their skills in data analysis and interpretation; the focus of others may be data stewardship; still others may be interested in software or algorithm development, and so on. The level at which training is required may also vary, according to whether trainees need a working knowledge of some tool or database to allow them to address routine tasks, or a deeper knowledge that would allow them to move the field forward. Depending on trainees’ objectives, the depth, length and organisation of the training offered will need to be adjusted to meet the desired outcomes. Sometimes, however, although individuals may know what it is they want to be able to do as a result of their training, they may not know which route offers the best pathway to achieve this.

### 2. Background knowledge

Previous experience and background knowledge also shape trainees’ needs, and the type of training likely to be most appropriate for them. For example, given the objective ‘to become competent using NGS data-analysis workflows’, the training required by wet-lab scientists with pure molecular biology backgrounds is likely to be rather different from that required by computational biologists, proficient at the command line and fluent in scripting. So again, for training to be effective, the nature of what is offered will need to be adjusted to suit trainees’ backgrounds.

### 3. Format of training

Finding the most suitable format for training (including depth, level of difficulty, interactivity, *etc.*) depends on a complex interplay between individuals’ reasons for seeking training, their background knowledge and experience, and how they want to learn. For example, if bioinformatics is to be a focus area for trainees, then it makes sense to invest time in learning generic programming and statistical skills; building on these foundations, they can then take more advanced courses (*e.g.*, generalised linear models, parallelisation in R), as necessary. Conversely, if the objective is point-of-need training, the imperative is to up-skill individuals for a specific task (*e.g.*, NGS data analysis) as efficiently as possible - here, if trainees have no experience with the command line or programming, it might be more appropriate to deliver the relevant analysis principles via an interactive Graphical User Interface (GUI). Their backgrounds and needs will hence determine how individuals want to learn, and will dictate the most appropriate training format and tools for them.

### 4. Time available for training

Clearly, the time available to receive training constrains the choice of training format. For example, a clinical researcher might not have time to become fluent in R coding, so may prefer a training programme that concentrates on analysis theory and uses analysis-tool GUIs. Conversely, a dedicated bioinformatician is more likely to want to invest time in becoming proficient with command-line tools and programming. Inevitably, the time needed for training depends on the level of skill that trainees desire to achieve. There is therefore often a tension between the amount of time trainees are able to devote to training and the time it realistically takes to gain the skills they need. This can lead trainees to have unrealistic expectations of what can be achieved in the allotted time, which, in turn, requires careful management of their expectations.

Although much of the above may seem self-evident, it is important to realise that these and other factors are likely to have influenced the survey responses. For example, the time available for training will probably have had a role in determining how individuals have received bioinformatics training (whether focused and deep, broad and shallow, *etc*.); it will likely have shaped the level of confidence they have achieved in using bioinformatics databases, software and command-line tools; it probably dictates what skills they still need (whether foundational or niche), and how they would prefer to acquire bioinformatics training in the future, given their particular time constraints. Similarly, the background knowledge of individuals is likely to have determined their reasons for seeking training, how confident they are in using bioinformatics tools and resources, the skills they still need and their preferred format for future training.

## Take-home messages from the surveys

Mindful that the surveys cannot be considered rigorous or complete, the results nevertheless highlight some training gaps across the globe, and preferences for how bioinformatics training should be delivered in order to address those gaps. Just as it is important to consider factors that are likely to have influenced the responses, such as the nature of the cohort surveyed, it is also important to think about how the survey questions were phrased if we are to arrive at meaningful conclusions. For example, more than 75% of survey respondents were beyond MSc/PhD stage (Figure 1). Hence, when asked, “*How would you prefer training to be delivered to you?*”, these individuals are more likely to have recommended face-to-face courses/workshops and online resources because i) these formats are best able to deliver training at the point-of-need, and ii) at these later career stages, UG and PG courses are generally no longer the most relevant or practicable vehicles for learning. Phrased differently - *e.g.*, “*How should bioinformatics training be delivered?*” - the question might have elicited rather different answers. Not surprisingly, then, formal degree programmes emerged as the least popular routes for receiving future bioinformatics training, despite their obvious importance in providing fundamental bioinformatics education.

As we have seen, acquiring greater expertise and confidence in data analysis and interpretation were amongst the most important needs reported, short courses and workshops being highlighted as the most popular ways to help plug skills gaps. A number of specific suggestions were also made concerning training resources to make resources more helpful for learners. Examples include i) having access to a range of case studies; ii) the ability to join ‘study groups’ where beginners could receive greater encouragement and peer support when learning how to use bioinformatics software tools and databases; and iii) the ability to evaluate their skills after training (as is routine in traditional coursework), perhaps via some kind of accreditation mechanism, thereby formalising and standardising the skill gain from point-of-need training programmes.

Perhaps the most urgent, overriding message, however, was that the next generation of life scientists should be reminded of the importance of bioinformatics and biostatistics throughout their studies, and that the learning process should commence with basic courses at UG level, to introduce and instil necessary skills at an early stage - ultimately, that bioinformatics should be woven into the fabric of future life-science degree programmes.

## The challenge for bioinformatics training

The survey data suggested that most individuals were likely to seek bioinformatics training at the point-of-need, when data had been generated, presumably because relevant training either had not been received during their academic education or had not been specific enough to address the research problem at hand. Just-in-time training was therefore their preferred mode of learning bioinformatics, particularly via online, self-taught modules.

We feel that this is too late, not least because the foundational, broad-based bioinformatics education is not in place - the point-of-need training strategy only really works when basic elements are already there to build on. This view was echoed in some of the survey free-text responses, one of which stated rather emphatically, “*courses at institutes are complete failure because they are so short and shallow. I think short courses at institutes only help people that already have skills to apply these skills on specific problems. I think programming is an essential skill for any scientist and should be taught in schools!*”

Part of the problem is that the later the intervention, the more difficult it is to achieve trainee self-confidence and skill retention. We regularly observe repeat trainees who have simply forgotten what they learned previously, because they were not able to put the skills they had acquired into regular practice - the ability to perform a particular type of analysis or to write a script is quickly forgotten if these tasks are not performed routinely in the weeks and months following a given course or workshop.

Often, trainees prefer training at the point-of-need in order to save time. However, some then realise that their training should have started before they began designing their experiments and collecting the data they now need to analyse. It seems that the earlier computational thinking is introduced, and skills in handling anything from command-line interfaces to programming are embedded, the longer the retention and greater the sustained confidence we can achieve in trainees. So, while there is clearly a very strong demand for just-in-time training, in the longer term, investment in formal training at UG level (if not sooner) seems to be beneficial for actual skill gain and retention.

In his blog, Sean Eddy made a call for all “*biologists to learn to do their own data analysis*” and to pick up “*scripting as a fundamental lab skill, like pipetting*” (Eddy, 2014). While not all biologists need to (or can) learn programming, introducing bioinformatics earlier into the education cycle could help to re-balance the number of computationally-minded biologists who can populate wet-lab teams and manage the programming and statistical components of data analyses. This view was echoed in the workshop and in survey responses, one of which lamented, “*It is too late for me, we need to fix the undergraduate syllabus - all biology students should be encouraged to learn programming and statistics*”.

Given the global preference for point-of-need training, however, and the attendant skill-development and -retention issues, it may be worthwhile to revisit current short-course and online bioinformatics training programmes, and to segment larger bioinformatics analyses into smaller ‘bite-sized’ chunks for online delivery in an attempt to facilitate skill uptake and retention at the point-of-need. Regardless, and at the least, individuals seeking bioinformatics training could receive better guidance in choosing the best training approach and resources to meet their particular training needs.

This global bioinformatics training demand brings with it a need for more and larger cohorts of individuals capable of delivering just-in-time training on specific topics to local communities. Relying on existing academic tutors, even those with computational or statistical backgrounds, to establish point-of-need training programmes is not always successful, as they may have no experience in the required, niche-specific bioinformatics topics (*e.g.*, sequence alignment in RNA-seq for isoform discovery). Moreover, while skilled in delivering traditional didactic coursework for semestered academic courses, they may struggle to adjust their content to short training programmes if they have not received guidance on how best to do so (Via *et al.*, 2013). This situation argues for the development of programmes tailored for future trainers, to disseminate best practices, to empower new bands of individuals to assuage local training needs and to ensure that trainers’ skills are kept up to date.

## How GOBLET can contribute to the global bioinformatics training landscape

As outlined above, there are several gaps in current global bioinformatics training provision, ranging from when training begins to a shortage of skilled trainers. GOBLET has an important role to play in addressing some of these gaps through its portal (Corpas *et al.*, 2014), best practice guidelines, guidance in training choice and trainer-support activities.

One issue that emerged very strongly from the surveys and workshop discussions was that the way individuals acquire bioinformatics skills, or the way they want to acquire them, changes along their career paths, essentially in proportion to the amount of ‘spare’ time they have available (Figure 4). GOBLET has a role to play here in providing free training materials in niche, specialised bioinformatics topics, as is often required for point-of-need training. The GOBLET portal collects materials from its global members and contributors, and makes them freely accessible under a creative commons licence (www.mygoblet.org/training-portal).

**Figure 4:**
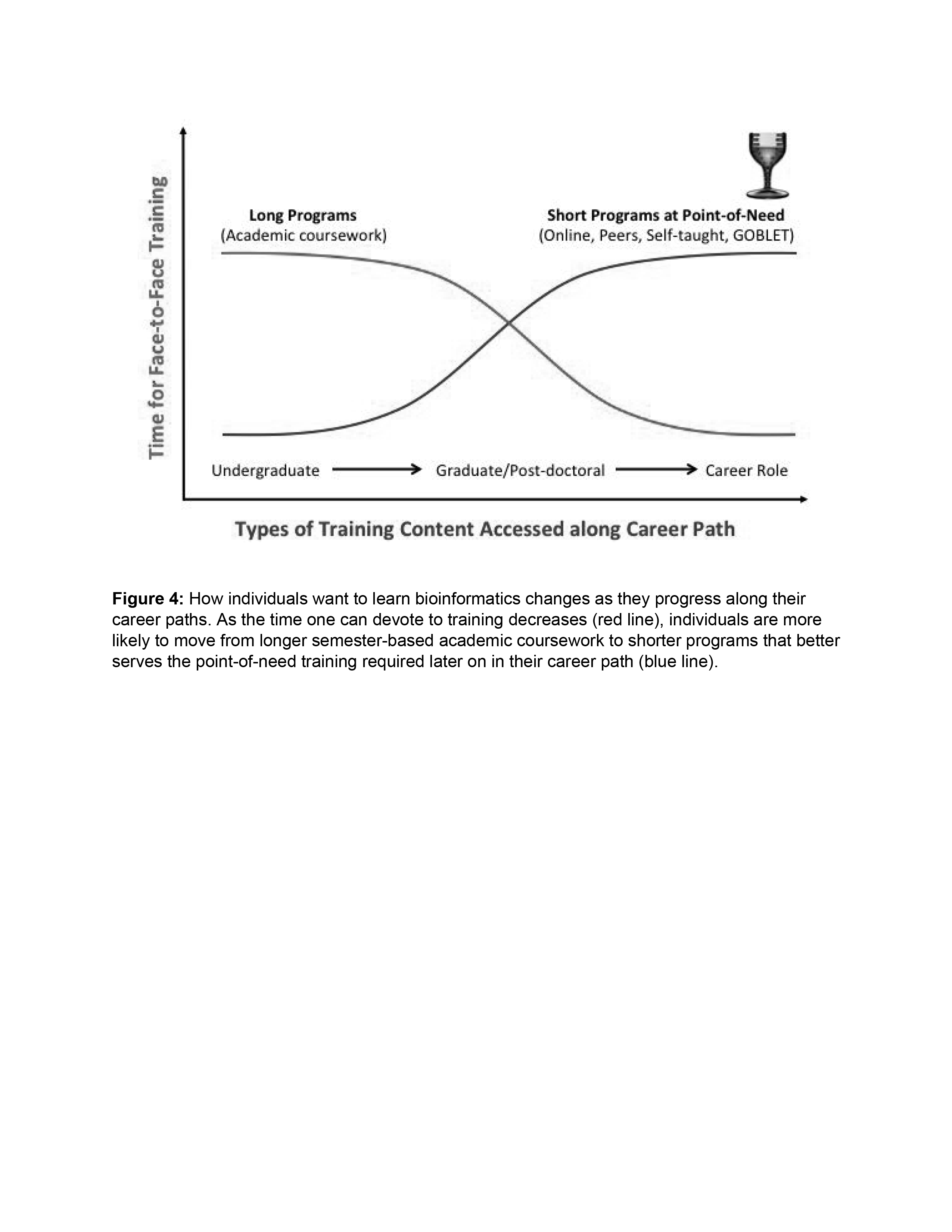
How individuals want to learn bioinformatics changes as they progress along their career paths. As the time one can devote to training decreases (red line), individuals are more likely to move from longer semester-based academic coursework to shorter programs that better serves the point-of-need training required later on in their career path (blue line).

Plans are underway to develop mechanisms for reviewing and evaluating materials and courses made available via the portal; we are also exploring recognition mechanisms for trainers (*e.g.*, via open badges). These will provide some measure of quality both for the training materials/courses and for trainers. Ultimately, by encouraging the use of standards, GOBLET aims to drive up the quality of *ad hoc* training courses.

To address the shortage of skilled point-of-need trainers, hand-in-hand with organisations like ELIXIR and H3ABionet, GOBLET is developing and delivering specialised workshops for new and more experienced trainers (Train-the-Trainer workshops), covering not only the specific bioinformatics topics to be taught, but also the principles important in providing robust training: how to define learning objectives, how to organise and set up short courses, how to develop and deliver materials, how to evaluate training results and so on. These best practices (Schneider *et al.*, 2010; Via *et al.*, 2013; Watson-Haigh *et al.*, 2013) are aspects of teaching that trainers might otherwise not be exposed to, but courses in which they are overlooked are less likely to satisfy their audiences: as one respondent commented, “*there is often a mismatch between the expectations/abilities of course attendees and the level at which a course is taught. People are generally only interested in learning about techniques that are directly applicable to their current research interests. On broader courses, this means they only benefit from a subset of the material that is available to them*”.

GOBLET’s Train-the-Trainer programme is developing a community of like-minded trainers with which to share experiences and materials; these materials are being regularly updated, as the field of bioinformatics evolves. This is important, as one survey respondent noted, because “*retraining the trainer concept/phenomenon must be a continuous thing in practice to adequately meet with the attendant progress made so far in bioinformatics*.”

Longer term, in light of these survey findings, GOBLET will consider conducting a bi-annual survey, both to capture bioinformatics training needs globally and to monitor how these needs are changing over time and jurisdiction. The ABPI survey, for example, viewed alongside its 2008 predecessor (ABPI, 2008), highlights that training needs do evolve over time; and the training deficit in computational aspects of biology may even widen as the data-science revolution takes hold, bringing with it new training imperatives (*e.g.*, in high-performance and parallel computing) for life-science students of tomorrow.

GOBLET will also consider establishing a decision tree to help guide trainees and trainers to clearly identify their bioinformatics needs, and to suggest what training steps are needed to meet these needs. Such a tool would be useful in: 1) identifying the right type of training for an individual, taking into consideration factors such as the training objective, trainee background knowledge, training format preference and time constraints, as outlined earlier; and 2) helping trainers to develop courses that match their audience needs. By providing better signposting, this may help to address a common frustration emerging from the surveys that there are, “*So many learning sites/tutorials/tools - [it is] difficult to pick the ‘best one*”.

GOBLET provides a community-based forum for discussing bioinformatics training issues, helping to identify and address real bioinformatics training challenges by running surveys, organising workshops, meetings, special interest groups, and so on. Results from our initial surveys have highlighted a number of important issues.

Respondents across the globe argued that universities should provide training and encouragement in the use of bioinformatics databases and software tools - above all, that they should provide more relevant training at an early stage. Until then, the demand for point-of-need training (whether face-to-face or online) in the analysis and interpretation of large volumes of complex, high-throughput life-science data will persist. By seeking the views of communities worldwide, and listening to their needs, GOBLET will continue to play an important role in helping to develop those programs, to enlarge the community of trainers and to plug current training gaps.

There is clearly still a lot of work to be done. Working alongside training groups like those within ELIXIR (with whom GOBLET has formulated a joint training strategy), the ISCB curriculum and competencies taskforce, CODATA-RDA, BD2K, the BBSRC Bioinformatics and Biomathematics Training Hub, and others that are emerging (“Canadian Bioinformatics and Computational Biology Strategic Framework,” 2015), GOBLET is now well placed to help build the infrastructure needed to quench the global thirst for bioinformatics training.

## Supplementary Data

**Supplementary Figure 1.**
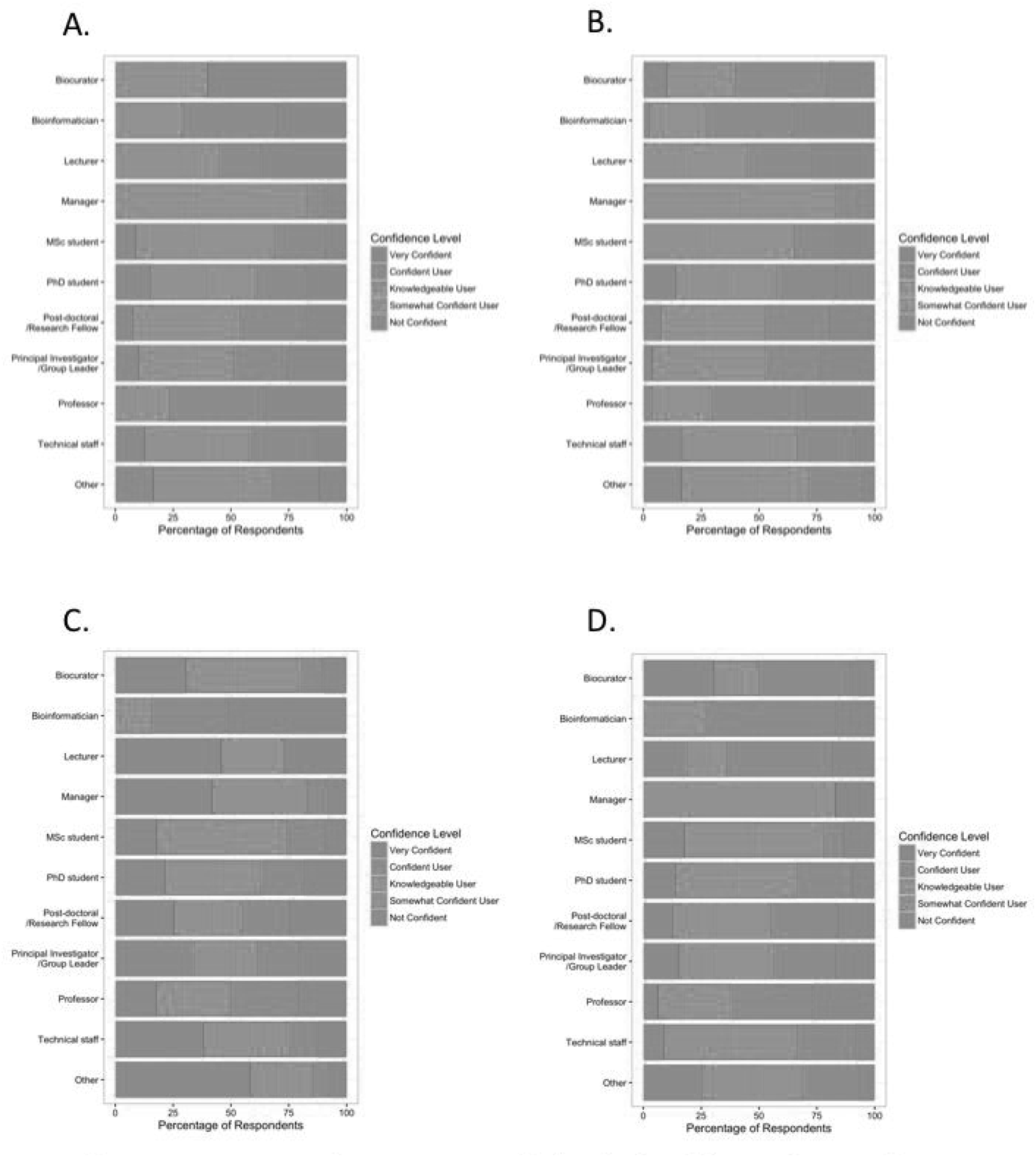
Charts comparing the level of confidence of respondents to the GOBLET survey in using bioinformatics databases (A), Web-based software (B), with command-line tools (C) and with data analysis and interpretation (D), on a 5-point scale from ‘Not confident’ (magenta) to ‘Very confident’ (red). Intervals on the x-axis denote the percentage of respondents for the given category.

